# A genome-scale study of metabolic complementation in endosymbiotic consortia: the case of the cedar aphid

**DOI:** 10.1101/179127

**Authors:** Miguel Ponce-de-Leon, Daniel Tamarit, Jorge Calle-Espinosa, Matteo Mori, Amparo Latorre, Francisco Montero, Juli Pereto

**Author notes:** Corresponding authors: - Miguel Ponce-de-Leon - Juli Pereto.

## Abstract

Bacterial endosymbionts and their insect hosts establish an intimate metabolic relationship. Bacteria offer a variety of essential nutrients to their hosts, whereas insect cells provide the necessary sources of matter and energy to their tiny metabolic allies. These nutritional complementations sustain themselves on a diversity of metabolite exchanges between the cell host and the reduced yet highly specialized bacterial metabolism –which, for instance, overproduces a small set of essential amino acids and vitamins. A well-known case of metabolic complementation is provided by the cedar aphid *Cinara cedri* that harbors two co-primary endosymbionts, *Buchnera aphidicola* BCc and *Ca.* Serratia symbiotica SCc, and in which some metabolic pathways are partitioned between different partners. Here we present a genome scale metabolic network (GEM) for the bacterial consortium from the cedar aphid *i*BSCc. The analysis of this GEM allows us the confirmation of cases of metabolic complementation previously described by genome analysis (*i.e*. tryptophan and biotin biosynthesis) and the proposal of a hitherto unnoticed event of metabolic pathway sharing between the two endosymbionts, namely the biosynthesis of tetrahydrofolate. *In silico* knock-out experiments with *i*BSCc showed that the consortium metabolism is a highly integrated yet fragile network. We also have explored the evolutionary pathways leading to the emergence of metabolic complementation between reduced metabolisms starting from individual, complete networks. Our results suggest that, during the establishment of metabolic complementation in endosymbionts, adaptive evolution is more significant than previously thought.

## Introduction

Species coexisting in a determined environment establish a network of interactions moulded by biotic and abiotic factors (Faust and Raes, 2012; Seth and Taga, 2014). From a molecular point of view, such networks can be considered as an entangled circuitry of various metabolisms interconnected by the exchange of compounds. In the mutualistic symbioses, where partners exchange nutrients or precursors bi-directionally, the nutritional interdependence will lead, in most cases, to a co-evolutionary process. A particular case occurs when the host cells harbour one or more symbionts inside them (*i.e.* endosymbionts). As a consequence of the adaptation to the intracellular life, the endosymbionts undergo many biochemical and structural changes, with extreme genome reduction by gene loss being the most dramatic one, compared to their closest free-living relatives (Manzano-Marín and Latorre, 2016; Moran, 1996; Moran and Bennett, 2014; Moya et al., 2008). Gene losses in endosymbionts result in the total or partial demolition of metabolic pathways, and thus endosymbionts become auxotroph for a diversity of compounds, such as nucleotides or amino acids.

The description of nutritional interactions between hosts and symbionts (and among symbionts in consortia) usually relies on the concept of “metabolic complementation”, for which at least two distinct meaning have been used. First, we can consider the exchange of essential components (*e.g*. vitamins and amino acids) between host and endosymbiont (López-Sánchez et al., 2009; Macdonald et al., 2012; Moya et al., 2008; Russell et al., 2013; Wu et al., 2006), a phenomenon usually known as cross-feeding. Second, metabolic complementation also refers to more complex scenarios where pathways can be fragmented and distributed between the members of the association (Price and Wilson, 2014; Van Leuven et al., 2014). Various examples of metabolic complementation in insect endosymbionts have been described in the past (Baumann et al., 2006; Manzano-Marín and Latorre, 2016; McCutcheon and Moran, 2011). As an example, some endosymbionts hosted by phloem-feeding insects (*e.g. Buchnera aphidicola, Candidatus* Tremblaya princeps, *Candidatus* Portiera aleyrodidarum) upgrade the host diet by supplying essential amino acids and vitamins absent in the diet (Baumann et al., 2006; McCutcheon and von Dohlen, 2011; Zientz et al., 2004). In other cases, as the cockroach endosymbiont *Blattabacterium cuenoti,* the assembly of metabolic reactions from the host and the symbiont allows the mobilisation of the host nitrogen reservoirs (Patiño-Navarrete et al., 2014).

A remarkable case of endosymbiont consortia has been described in the cedar aphid *Cinara cedri* (Lamelas et al., 2011a; Pérez-Brocal et al., 2006). In this system, two species of endosymbiotic bacteria coexist. As it is the case in most aphid species, the primary (obligate) endosymbiont is *B. aphidicola* BCc, albeit in this insect species there is always a second (co-primary) endosymbiont, *Candidatus* Serratia symbiotica SCc (hereafter referred to as *S. symbiotica* SCc, Gómez-Valero et al., 2004). The genomic analysis of this consortium has shown that many biosynthetic pathways are coded only in one of the two endosymbiont genomes, thus leading to obligate cross-feeding. Nonetheless, it is remarkable that the tryptophan biosynthetic pathway is split in two halves: *Buchnera* is able to synthesize up to anthranilate, whereas *Serratia* uses this anthranilate to synthesize tryptophan, which is required by all the members of the consortium (Gosalbes et al., 2008). The existence of metabolic complementations between endosymbionts and their host and, particularly, between the members of the bacterial consortium (as it is the case with the biosynthesis of tryptophan), poses several evolutionary questions: does complementation generate an adaptive advantage for the system as a whole? And if this is the case, what is the nature of such advantage? Some theoretical studies have tried to illuminate whether the organisms exhibiting some degree of cooperation, as it is the case of cross-feeding, show some increase in their growth rate, compared to non-cooperative strains (Germerodt et al., 2016; Großkopf and Soyer, 2016). Furthermore, the metabolic pathway sharing between two endosymbionts has been suggested as a strategy to increase the efficiency of the biosynthesis of compounds when feedback inhibition is present in the pathway (Mori et al., 2016). Nevertheless, the emergence of such patterns of complementation poses diverse biophysical problems. For instance, the exchange of intermediates between endosymbionts implies a transport of solutes. It is well known that endosymbionts harbour a limited repertoire of transporters (Charles et al., 2011) and, on the other hand, the existence of membrane transporters specific for metabolic intermediates is very unusual. As a result, the exchanges should occur by simple diffusion and this situation imposes restrictions since metabolic intermediates usually show very low diffusion rates (Mori et al., 2016).

Another important matter in the evolution of endosymbiotic bacteria is the interplay between chance and necessity during the genome reduction process (Sabater-Muñoz et al., 2017). Although it is reasonable to accept the force of purifying selection, it is not clear if the patterns of complementation exhibited in these systems are the outcome of a random process, or if the observed patterns reflect an advantage over alternative evolutionary trajectories. In other words, to what extent are the evolutionary histories of these systems predictable? *In-silico* evolutionary experiments using genome-scale metabolic models (GEM) of two different endosymbionts (*B. aphidicola* and *Wigglesworthia glossinidia*) showed that the present gene content of these symbionts can be predicted with over 80% accuracy, from distant ancestors of the organisms and considering their current lifestyle (Pál et al., 2006). Similar studies have analysed the fragility of the reduced metabolism to conclude that, in general, these networks cannot be further reduced (Belda et al., 2012; Calle-Espinosa et al., 2016; González-Domenech et al., 2012; Pál et al., 2006; Ponce-de-Leon et al., 2013; Thomas et al., 2009). A recent application of metabolic flux analysis to an endosymbiotic consortium has revealed distinct benefits and costs of the symbionts to their host (Ankrah et al., 2017), highlighting how the analysis of GEMs can be successfully applied to study endosymbiotic consortia.

Herein, we are interested in studying the interplay of chance and necessity in the evolution and emergence of metabolic complementation in endosymbiotic consortia. For this purpose, we have chosen the well documented case of the consortium formed by the primary and co-primary endosymbionts of the cedar aphid, *B. aphidicola* BCc and *S. symbiotica* SCc. Individual GEMs have been reconstructed, manually curated and analysed for each individual bacterium, based on their corresponding genome annotations. After extensive manual curation, these two models were used to create a compartmentalized consortium model named *i*BSCc, which also include a set of key enzymatic activities performed by the host. We assessed the metabolic connections between the two endosymbionts and also with the host to predict patterns of metabolic complementations. We observed that the combination of these two extremely reduced metabolic networks results in an integrated yet fragile system. Finally, we performed *in-silico* evolutionary experiments to study the paths leading to the emergence of metabolic complementation.

## Materials and Methods

### Annotated genomes

In order to reconstruct the genome-scale metabolic models (GEM) of *S. symbiotica* SCc and *B. aphidicola* BCc, we retrieved the corresponding genomes and the semi-automatically reconstructed Pathway-Genome Databases (PGDB) available in the BioCyc collection in (Caspi et al., 2014). The used PGDBs’ versions were SSYM568817 19.0 for *S. symbiotica* SCc, and BAPH37261 19.0 for *B. aphidicola* BCc (both available in Tier 3 at BioCyc 19.0) (See supplementary Text S2). The public version of BAPH372461 does not include the information encoded in the pTpr-BCc plasmid (accession number EU660486.1 (Gosalbes et al., 2008). This plasmid contains two genes (*trpE* and *trpG*) (Table 1), and was manually added to a local version of the *B. aphidicola* BCc PGDB, created through Pathway Tools v. 19.0 (Karp et al., 2015). Once added, the PathoLogic algorithm was run again in order to update the pathway prediction.

**Table 1.**
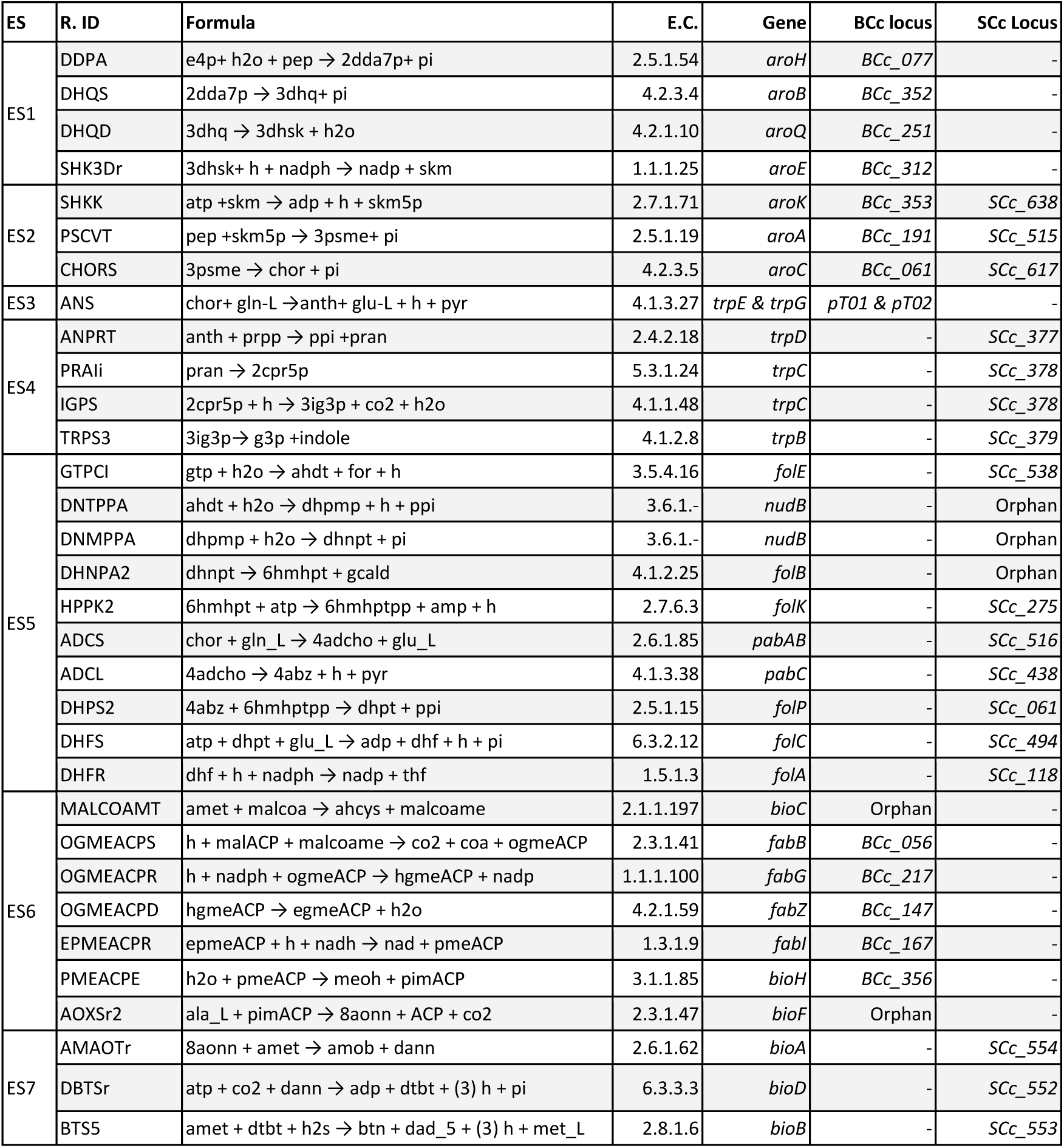
Enzymatic activities and genes involved in the biosynthetic pathways of tryptophan, THF and biotin as grouped in 7 enzyme subsets (ES). The IDs for the reactions and the metabolites are as obtained from the BiGG database.

### Reconstruction and refinement of the metabolic models

The reconstruction and refinement of the metabolic models was performed following the protocol described by Thiele and Palsson (2010). In order to reconstruct the metabolism of *S. symbiotica* SCc, the GEM of *E. coli* K12 MG1655 *i*JO1366 (Orth et al., 2011) was used as a reference, since this is the phylogenetically closest free-living organism for which a highly refined and validated model exists. First, orthologous genes were identified between the *E. coli* K12 MG1655 genome (available in EcoCyc 19.0) and the *S. symbiotica* SCc genome included in the PGDB SSYM568817 19.0. Gene sequences were extracted, translated and compared using *Blastp* (Altschul et al., 1990) with an e-value maximum of 10_-10_ and an identity minimum of 75%. Orthologs were then identified using OrthoMCL (Li et al., 2003). Combining these results with the set of reactions and pathways present in the SSYM568817 PGDB, a gene-protein-reaction (GPR) table was constructed. Using this GPR table, together with the *i*JO1366 model and the BiGG database (Schellenberger et al., 2010) a first version of the *S. symbiotica* SCc model was reconstructed. Moreover, the biomass equation introduced in the model is a modified version of the one present in *i*JO1366, from which we removed membrane components and cofactors absent in the *S. symbiotica* SCc network. The coefficients were corrected using the methodology suggested by Henry et al. (2010). The first draft obtained was revised combining the unconnected module (UM) approach (Ponce-de-Leon et al., 2013) together with the previously published genome analyses (Lamelas et al., 2011a). Regarding the metabolism of *B. aphidicola* BCc, a previously published GEM, named BCc (Belda et al., 2012), was used as a reference. However, this GEM contained several blocked reactions and dead-end metabolites. The resolution of the different gaps was performed by solving the set of UMs combining the pathway inferences present in the BAPH372461 PGDB and the GEM of *B. aphidicola* Bap (MacDonald et al., 2011), available in BioModels (MODEL1012300000).

### Construction of a biomass equation for the aphid *C. cedri*

The aphid biomass equation was defined by including the various biomass components in corresponding stoichiometric proportions. Since the consortium model does not include the metabolism of the host, the aphid biomass equation only includes the compounds that are provided by the endosymbiotic consortium (*i.e.* the essential amino acids, vitamins and cofactors) as previously done (Calle-Espinosa et al., 2016). The stoichiometric coefficients for the set of amino acids provided by the endosymbionts were estimated from the analysis of composition of the *Aphis fabae* and *Acyrtosiphon pisum* proteomes (Douglas et al., 2001; Russell et al., 2014). The compositions measured in two different aphid species showed good agreement, a fact that suggests that the values can be extrapolated to *C. cedri* (see Supplementary Text S1). We used the values from the experimental measurements of the *A. pisum* amino acid composition obtained by Russell et al., (2014) because this dataset includes the measurements of certain amino acids not included in Douglas et al. (2001). The values were normalised to represent composition in 1 g of dry weight (DW) in the aphid biomass. Additionally, the cofactors provided by the endosymbiotic consortium to the host were added. Since there is no estimation of their proportion in the aphid biomass, their values were set to be several orders of magnitude below the amino acids demand, but reflecting their essentiality to the host (Supplementary Table S5).

### Constraint-based modeling methods

The different constraint-based methods used in the present work correspond to the current implementation found in COBRApy toolbox 0.6 (Ebrahim et al., 2013). The boundary conditions used in the individual analyses of the endosymbionts models, *i*SCc226 and *i*BCc98, can be found in Supplementary Tables S1 and S2, respectively. However, in the case of the compartmentalized model of the consortium, a restriction that limits the total number of carbon atoms diffusing through the membrane of each endosymbiont was introduced (Burgard et al., 2001). By doing so, we are able to separately constrain the exchange of metabolites of each endosymbiont with its surrounding environment without the need of establishing arbitrary boundary for each exchange flux. Thus, instead of having a large set of parameters, one for each exchange flux, we only used two parameters, one for each endosymbiont. The restriction can be expressed as the following linear inequality:

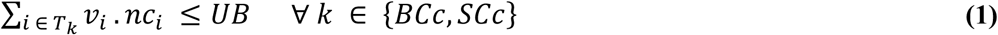

Where *T*_*k*_ *⊂ J* is the set of indexes of the transporters present in the compartment ***k*** (endosymbiont) and *J* the set of all fluxes; *v*_*i*_ is the flux through the transporter *i*; *nc*_*i*_ is the number of carbon atoms in the transported molecule; and *UB* is the upper bound, *i.e.* the parameter establishing the maximum number of carbon atoms that can be exchanged by the α compartment (one of the endosymbionts) and the extracellular compartment (the host cell). This parameter limits the amount of matter flowing through the membrane in terms of the total carbon atoms and was fixed to 100 carbon atoms. In order to guarantee that the transport fluxes are non-negative, the reversible transporters were split into two irreversible transport reactions with opposite direction (Burgard et al., 2001). The biosynthetic capabilities of the endosymbionts for each biomass component, as well as for energy production, were assessed using Flux Balance Analysis (FBA) (Orth et al., 2010). Furthermore, network fragility was predicted through in-silico knockout experiments conducted using FBA as well as the Minimization of the Metabolic Adjustment (MOMA) (Segrè et al., 2002). Details on each method can be found in the extended materials and methods (see Supplementary Text S2).

## Results

### Metabolic reconstruction of the cedar aphid primary and co-primary endosymbionts

The metabolic models of *S. symbiotica* SCc and *B. aphidicola* BCc were reconstructed individually, and were named *i*SCc236 and *i*BCc98, respectively (see Supplementary Text S1 for further details on the reconstruction, and Supplementary Tables S1 and S2 for the complete models). The former consisted of 267 intracellular metabolites and 209 reactions catalysed by the products of 236 genes, plus 11 orphan reactions (*i.e.* reactions with unknown coding genes). It also includes 30 transporters associated with a gene, and 49 orphan transport reactions. On the other hand, *i*BCc98 yielded a smaller network, containing 155 intracellular metabolites and 95 reactions catalysed by the products of 98 genes, and 8 additional orphan reactions. Additionally, it includes only 5 transporters associated with a gene, and 58 orphan transporters. In both cases, all orphan reactions are required by the model in order to predict biomass formation. The genomes of these endosymbionts contain only a small set of genes coding for substrate-specific transport systems (Charles et al., 2011; Wernegreen, 2002). However, the corresponding metabolic models predict the necessity of metabolite transit through the endosymbiont membrane, albeit the transport mechanisms have not been elucidated in many cases, in which simple diffusion has been proposed as a plausible mechanism (Mori et al., 2016)The metabolic requirements and the biosynthetic capabilities for each of these two models were congruent with those inferred from genomic analyses (Lamelas et al., 2011a), with the exception of the biosynthesis of asparagine, which is predicted by the *i*SCc236 model to be required as nutritional input provided by the host instead of being synthesized by *S. symbiotica* SCc (Supplementary Fig. S1). Finally, when the energetic capabilities, *i.e.* the synthesis of ATP, was analysed for both symbionts, we found that *i*BCc98 predicts a very low yield of ATP, limitation that is a direct consequence of the absence of ATP synthase. In turn, this lack of a proton pump mechanism poses a constraint on the regeneration of NADH, through the NADH dehydrogenase complex. The model suggests that part of the NADH may be driven through the conversion of the pair homocysteine and serine, into glycine and methionine (for further details, see Supplementary Text S1).

Since we were interested in modeling the whole consortium, the two metabolic models previously introduced (*i*BCc98 and *i*SCc236), were combined to create a single model named *i*BSCc. Although it is known that *B. aphidicola* BCc population and *S. symbiotica* SCc are hosted in different bacteriocytes, a more simplistic representation was chosen where both endosymbiont models are embedded in a single compartment, in a similar way as it has been previously done (Ankrah et al., 2017). Therefore, our model included: i) a compartment representing the *B. aphidicola* BCc population; ii) a compartment representing the *S. symbiotica* SCc population; and iii) a single extracellular compartment representing the host cells, where both symbionts are embedded (Fig. 1). Consequently, the boundary of the system was defined by the compartment representing the host, and exchange fluxes across the boundary represented the metabolites supplied and consumed by the aphid, as well as the excretion products. The 52 metabolites found in the extracellular compartment include those obtained from the host, common excretion products and metabolites exchanged by the two symbionts. Additionally, five reactions were added to the extracellular compartment, as they have been suggested to be performed by the host and to play a relevant role in the metabolic complementation between host and symbionts (Hansen and Moran, 2011; Poliakov et al., 2011). These reactions include the conversion of phenylalanine into tyrosine (1 reaction), the production of homocysteine and adenosyl-methionine from cysteine (3 reactions) and the assimilation of sulphydric acid, produced by S. symbiotica SCc, for the production of lipoate (1 reaction) (see Supplementary table S3).

**Figure 1.**
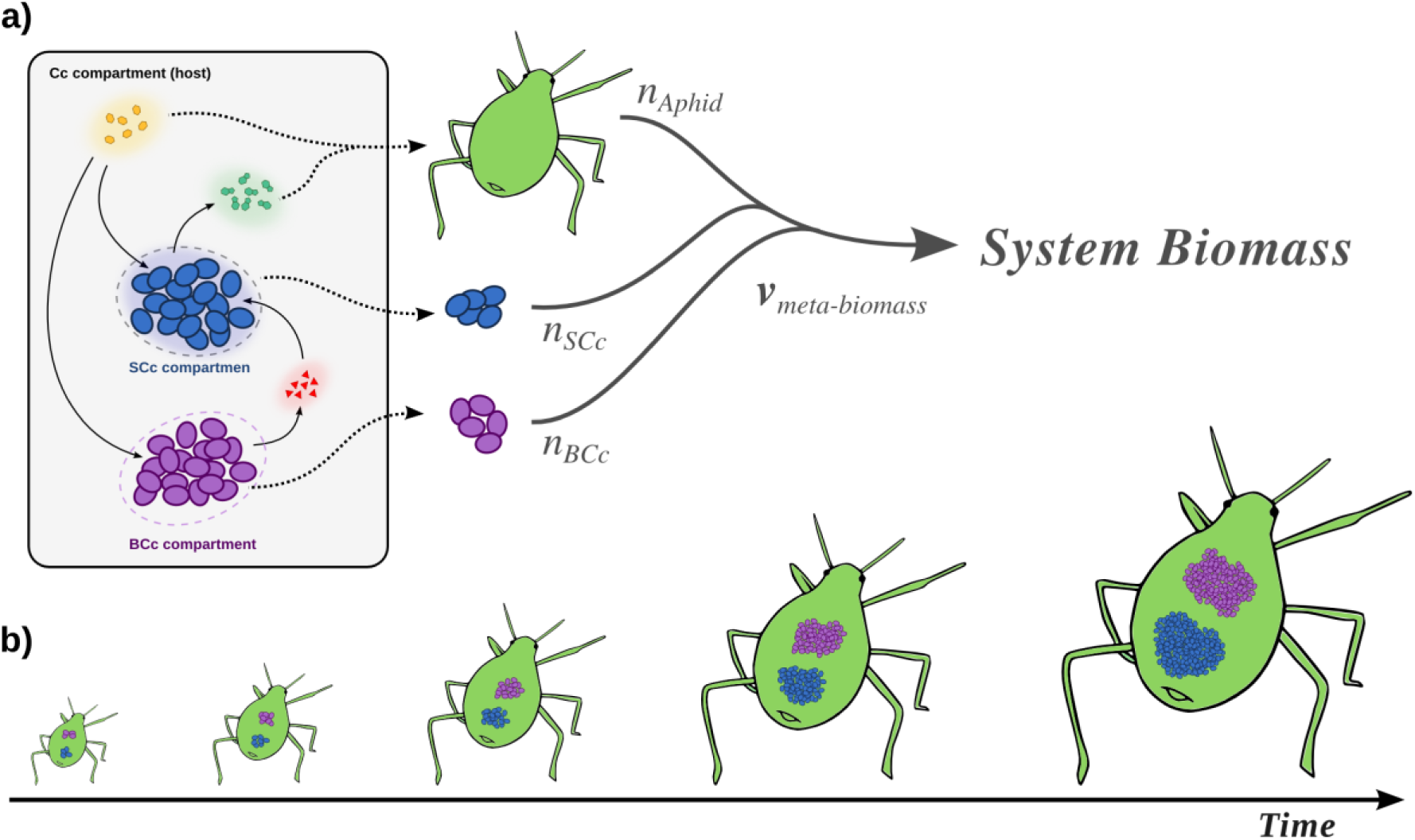
Compartment modeling of the cedar aphid endosymbiotic consortium and meta-biomass equation modeling the growth of the endosymbiotic consortium and the host. a) Representation of the three compartments model of the endosymbiotic consortium. Dotted arrows represent the biomass production of each member. The thick tripartite arrow represents the system biomass, i.e. the meta-biomass (see main text). The annotations nα with α ∈ {BCc,SCc,Aphid} correspond to the stoichiometric contribution of each member to the total system biomass equation. Solid arrows: transport processes within the system (e.g. metabolic complementation). b) Coupling between the growths of the members in the system, representing stability in the ratio of their biomass.

In order to simulate the growth of the system as a whole, we introduced a combined meta-biomass equation, where each member contributes to the growth of the system with a fixed stoichiometry (Fig. 1a). Our model assumes the coupling between the growths of all the members of the system, which would especially apply during the development of the host (Fig. 1b). This assumption is justified by different sources of experimental data as well certain theoretical results (see Supplementary Text S1 for further details). Since the exact contribution of the symbionts to the cedar aphid biomass is unknown, we used data obtained from *Schizaphis graminum* and *A. pisum* indicating that their symbionts represent 5-15% of the system’s biomass (Baumann et al., 2006; Whitehead and Douglas, 1993). Thus, we modelled the proportion between biomass of the symbionts and the host to be 1:9. Finally, since imaging data from the cedar aphid bacteriome indicates similar proportions between the two symbiont species (Gomez-Valero et al., 2004; Pérez-Brocal et al., 2006), we modelled that each bacterial member represents 5% of the system’s biomass at every time. It is important to emphasize that this assumption is key for the direct utilization of FBA and other related techniques in the study of the *i*BSCc model. The reason is that, if both symbionts are in proportion 1:1, the fluxes are normalized by the same quantity (e.g. dry weight of bacteria) and they grow necessarily at the same growth rate. This means that they behave essentially as one “big” bacteria. However, if this assumption does not hold, and the growth rate of the two bacteria are the same (i.e. they are coupled), new (probably non-linear) constraints should be incorporated to the model (see Kerner et al., 2012 as an example). Once the meta-biomass equation was introduced, we use FBA to verify that the *i*BSCc model is consistent, allowing for a positive meta-biomass flux and the growth of each member of the consortium (see Supplementary Tables S3). This optimal flux distribution predicted by FBA (maximization of the meta-biomass, *i.e.* the growth rate of the *i*BSCc consortium) is analysed in the next section.

### Metabolic analysis of the cedar aphid consortium

The overall structure of the models corroborates that the metabolic network of *B. aphidicola* BCc is specialised in the production of amino acids, while *S. symbiotica* SCc produces nucleotides and a large number of cofactors (Supplementary Fig S1). Moreover, the model predicts that *S. symbiotica* SCc can synthesize tryptophan from anthranilate, which has been shown to be provided by *B. aphidicola* BCc in a paradigmatic case of metabolic complementation (Gosalbes et al., 2008; Lamelas et al., 2011a; Manzano-Marín et al., 2016; Martínez-Cano et al., 2015). Another case of metabolic complementation between the two symbionts predicted by the model occurs in the biotin synthesis pathway, which takes place via the import of the precursor 8-amino-7-oxononanoate (8AONN), produced by *B. aphidicola* BCc, as recently suggested from genomic data (Manzano-Marín et al., 2016). Lysine biosynthesis also represents a case of complementation, based on the fact that the genome of *S. symbiotica* SCc encodes all activities of the pathway except the last one, which converts *meso*-diaminopimelate into lysine (Lamelas et al., 2011a). However, our model indicates that *meso*-diaminopimelate is used for the synthesis of peptidoglycan, while the complete lysine synthesis pathway is conserved in *B. aphidicola* BCc, suggesting that this complementation does not occur.

The FBA predictions showed that *B. aphidicola* BCc synthesizes and provides the host and *S. symbiotica* SCc with ten amino acids. *B. aphidicola* BCc also synthesizes anthranilate, 8AONN and shikimate and release them to the host compartment. In turn, *S. symbiotica* SCc imports anthranilate, 8AONN and shikimate, and use them as precursors for the biosynthesis of tryptophan, biotin and tetrahydrofolate (THF), respectively (see Fig 2 and Supplementary Table S3). Therefore, three events of metabolic complementation between the two symbionts are predicted, including the biosynthesis of tryptophan, biotin and THF; whereas the former two have been inferred previously from genomic analyses (Gosalbes et al., 2008; Manzano-Marín et al., 2016), the latter is a plausible as yet unidentified complementation case (see Fig 2). Folate cross-feeding has been described between *Serratia grimesii* and *Treponema primitia*, both members of the termite *Zootermopsis angusticollis* gut microbiome (Graber and Breznak, 2005). Since *C. cedri*, as animals in general, is not able to synthesize folate, and this is not included in the aphid’s diet, *Serratia* probably provides this essential cofactor to both the host and *Buchnera*.

**Figure 2.**
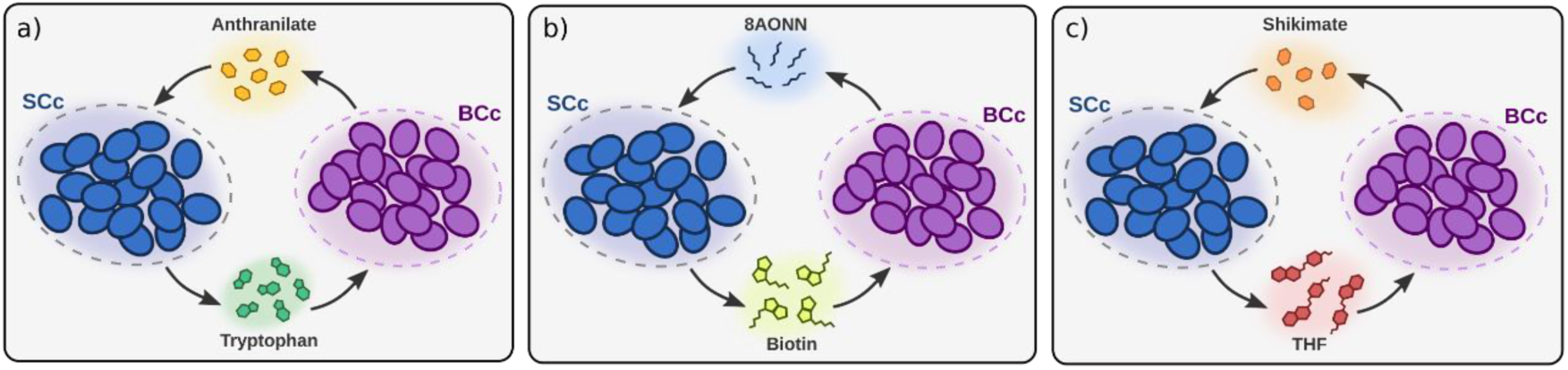
Patterns of metabolic complementation predicted by iBSCc. In a) the complementation for tryptophan biosynthesis, where BCc produces anthranilate and SCc uses it to produce tryptophan is shown. In b) and c) the complementation for the biosynthesis of biotin and THF, respectively, are shown.

Regarding *S. symbiotica* SCc, the *i*BSCc model predicts that it can synthesize, without need for complementation, cysteine, the four deoxynucleotides, the four triphosphate nucleosides, and thirteen cofactors and coenzymes. Several of these compounds may also be synthesized by the host (e.g. nucleotides, deoxynucleotides and NAD_+_), who might provide them to *B. aphidicola* BCc. Indeed, the aphid *A. pisum*, as most eukaryotes, is able to synthesize nitrogenous bases (Richards et al., 2010; Vellozo et al., 2011), which indicates that *C. cedri* also should. In this study, however, we will assume that it is *S. symbiotica* SCc who provides *Buchnera* with nucleotides and cofactors, and the host with cofactors such as biotin, riboflavin and THF.

### Fragility analysis: can the networks be further reduced?

The construction of the metabolic models of two different endosymbionts with a notable difference in size allowed us to study whether the reductive evolutionary trends indeed generate smaller, more fragile networks. The robustness of the metabolic networks of the cedar aphid endosymbionts was assessed through *in-silico* knockout analyses conducted with two alternative approaches: FBA and MOMA. In the first place, the fragility of the whole consortium was considered by using the meta-biomass flux as an indicator of the viability. The results show that around the 85% (∼88% using MOMA) of the genes coded by the endosymbionts are essential in order to sustain the growth of the whole system (see Supplementary Table S4). Then we focussed on the fragility of the individual endosymbionts networks. For the case of *i*BCc98 FBA predicted that 72 out of the 98 metabolic genes (∼74%) are essential, while the MOMA analyses identify 76 (∼78%) as essential. The dispensable genes are mostly involved in catabolism, affecting the phosphate pentose pathway, glycolysis, respiratory chain and pyruvate fermentation (see Supplementary Table S4). When performing FBA robustness analyses on *i*SCc236, 209 genes (∼88%) are predicted to be essential, whereas MOMA predicted 5 additional genes as essentials. The 28 dispensable genes predicted by both methods code for 36 enzymatic reactions involved mostly in biosynthetic pathways (*e.g.* nucleotides and cofactors), but also in the central carbon metabolism (*e.g.* the pentose phosphate pathway and glycolysis). If we consider the cell wall genes to be dispensable (since they have been repeatedly lost in endosymbiotic bacteria), the number of essential genes drops to 189 (∼80%). Additionally, if it is also assumed that the host is who provides the nucleotides and deoxynucleotides, the percentage of essential genes drops to ∼70% (data not shown). Although it might seem surprising that these estimates are higher or comparable to those obtained by using the smaller *i*BCc98 network, the *i*SCc236 model requires 22 organic compounds to be imported, while *i*BCc98 requires 29, among which there are nucleotides and cofactors such as NAD_+_ and coenzyme A. Finally, a study of distributed robustness was performed through the analysis of the synthetic lethal (or double lethal) genes, *i.e.* pairs of non-essential genes whose simultaneous inactivation yields lethality (Wagner, 2005). In the case of *i*BCc98, *ca*. 15% of the pairwise combinations between the 26 non-essential genes predicted with FBA are predicted as synthetic lethal. These combinations include 20 of the nonessential genes, indicating that most of the dispensable genes have only a shallow degree of redundancy. On the other hand, *i*SCc236 predicts only *ca*. 5% of the possible combinations between the 34 non-essential genes to be lethal. Moreover, by disregarding the cell wall biosynthesis genes as explained above, this number drops to *ca*. 2%.

### In-silico reduction experiments: evaluating alternative evolutionary scenarios

Although it is generally accepted that these patterns of complementation are the outcome of the process of genome reduction, whether such organization of the metabolic networks confer a selective advantage to the whole system or not, remains an open question. Previous work has focused on kinetic aspect of the problem, in particular the role of product inhibition as a plausible condition that may drive the emergence of metabolic complementation (Mori et al., 2016). Herein, we try to approach this problem from a structural point of view, by comparing the metabolic capabilities in alternative scenario of gene losses and retention within the three shared pathways. For this purpose, *i*BSCc was extended to represent a putative ancestral-state model of the consortium, named *i*BSCc_Ancest_, where the three shared pathways are still complete in both endosymbionts (see Supplementary Text S2). In this way, it is possible to compare how the different scenarios of gene loss and retention perform with respect to the putative ancestor as well as to the pattern of complementation exhibited by the cedar aphid consortium.

Using *i*BSCc_Ancest_ the space of all viable and scenarios of gene loss and retention patterns (GLRP) were generated by removing, from this model every possible combination of genes, from single genes to the most reduced cases where only one copy of each gene remains present (at least one of the two copies for each gene needs to be functional). A GLRP is considered viable if, after removing the corresponding reactions, FBA predicts a meta-biomass flux greater than zero. Furthermore, in order to reduce the number of combination of possible GLRPs, the enzyme subsets (ES) *i.e.* groups of enzymes that always work together under steady state, were first computed for *i*BSCc_Ancest_ (see the extended Material and Methods section in the Supplementary Text S2). Since removing an enzyme from an ES is equivalent to remove all the enzymes in the ES, each ES can be treated as a functional unit. Then, those genes coding for enzymes in the same ES were grouped together. Table 1 shows the structure of the ES for the three pathways considered in this study. Notably, the 32 enzyme activities are grouped in a total of 7 ESs. Due that in *i*BSCc_Ancest_ each endosymbiont includes the 7 ES, the enumeration process yield a total of 2188 viable GLRPs, which represent alternative consortium models. From this set, 128 are minimal GLRPs, i.e. only one copy of each gene remains active. It is worth to note that the pattern exhibited by the *C. cedri* consortium is not minimal since the genes coding for the three activities which allow the conversion of shikimate into chorismate are present (see Table 1). In order to simplify the notation, each GLRP was coded by a sequence of symbols 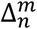, where the subscript *n* (from 1 to 7) denote the loss of the enzyme subset, by one of the endosymbionts, indicated by the supra index *m*, which can be either *B* or *S* (*B. aphidicola* or *S. symbiotica*). Accordingly, 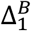 represents a simple GLRP where *B. aphidicola* has lost the ES1. A more complex example would be the case of *i*BSCc (which is the actual pattern exhibited by *B. aphidicola* and *S. symbiotica* in the cedar aphid). In this case, the GLRP is coded as follows: 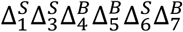. Note that in this case the ES2 is omitted because the genes involved in this ES are present in both endosymbionts, as mentioned above (see Table 1).

The 2188 consortium models, which represent alternative evolutionary scenarios, were evaluated using FBA in two ways by considering: i) the individual maximal production rate of tryptophan 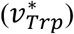 THF 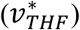 and biotin 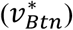; and ii the whole system performance, calculated by optimizing the meta-biomass production rate. In all the cases the optimal values are normalized with respect to the optimal value showed by *i*BSCc_Anc_. Figure 3 summarizes the results of the reduction experiment in terms of the production capabilities of tryptophan, THF and biotin for each GLRP. Firstly, the results show that for any of the three objectives, the production rates of the different GLRPs exhibit a great variability (for more details see Supplementary Table S5). This clearly shows that the way in which a pathway is distributed in a complementation event has a profound impact in the pathway capabilities. In the case of the tryptophan production, the different GLRPs can be divided into two main groups: i) a group of GLRPs (which includes *i*BSCc) with a tryptophan production rate almost equal to the one exhibited by the putative ancestor (*i.e.*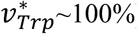); and ii) a larger group of GLRPs with production rate 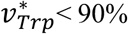< 90% (Fig 3a). Furthermore, when considering only the minimal GLRPs (*i.e.* cases with 50% of genes lost) the gap is even larger, and there is only one GLRP with 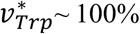 whereas the in other patterns 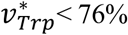. This minimal GLRP with 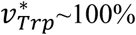 corresponds to the pattern exhibited by *i*BSCc with the additional loss of the only set of redundant genes that remain present which form the ES2 (see Table 1).

**Figure 3.**
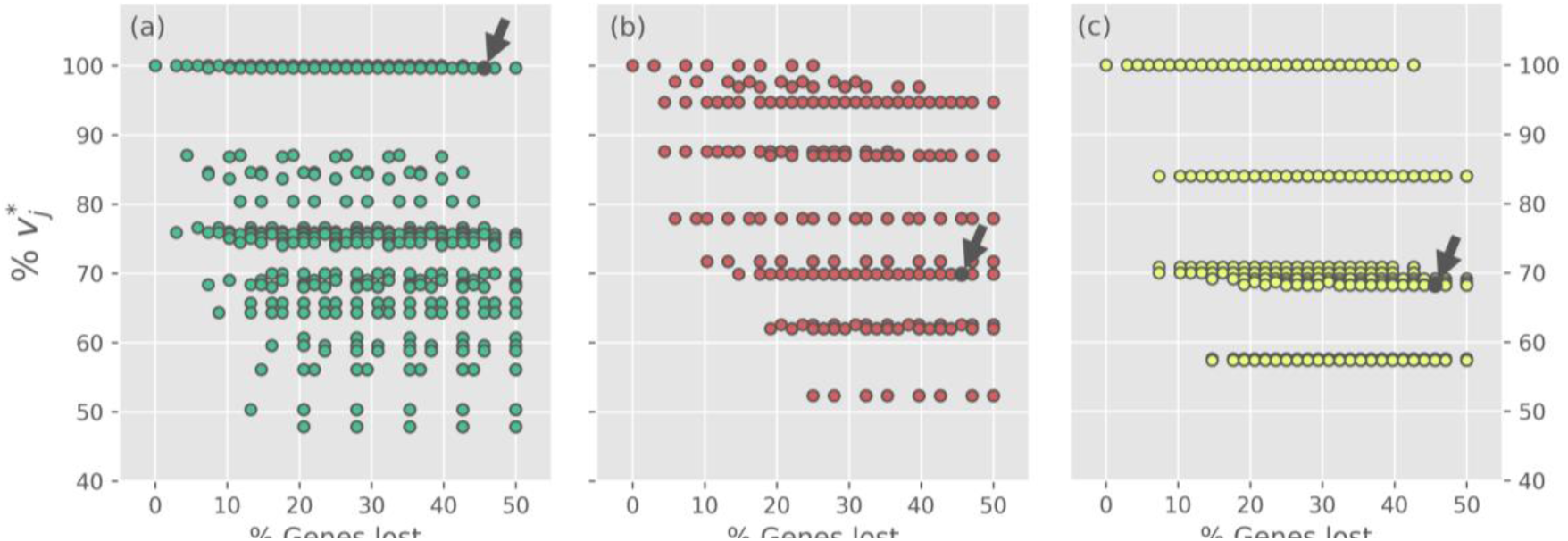
Optimal production rates of tryptophan, THF and biotin for the different genes loss and retention scenarios. In each panel the normalized optimal production rate 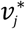*is plotted against the percentage of gene losses, for every gene loss scenario. From (a) to (c) the panels correspond to the optimization of the rate* 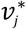 of production of tryptophan, THF and biotin, respectively (i.e.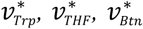). For each target, optimal production rates are normalized with respect the optimal value exhibited by iBSCc_Ancest_. The small arrow denotes the case of iBSCc, i.e. the cedar aphid consortium.

On the other hand, when considering the biosynthesis of THF and biotin, the results also show a wide dispersion for the normalized production rate values exhibited by the different GLRPs. However, unlike the case of the tryptophan, in these two cases the optimal production value exhibited by *i*BSCc decreases considerably with respect to the value exhibited by the putative ancestor (Fig 3b and 3c). For the case of the production of THF, all the GLRPs with the highest normalized production rate 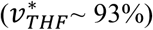 and with more than 40% of the genes lost, shared the following two losses: 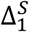 and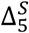, which correspond to the scenario in where *S. symbiotica* losses the biosynthetic pathways of shikimate and THF. Whereas, 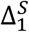 is consistent with the GLRP shown by *i*BSCc, 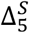 is the opposite, *i.e.* in the cedar aphid consortium *B. aphidicola* has lost the THF biosynthetic pathway. Furthermore, in all those GLRPs which involve 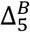 implies an important drop in the normalized production rate 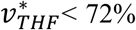 (Fig 3b). Something similar is found in the case of the biotin biosynthesis where the GLRP of *i*BSCc implies 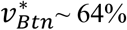 (Fig 3c). According to the results, when the system has lost more than 40% of the genes involved in the analyzed pathways, those GLRPs with a biotin production rate closest to the one exhibited by putative ancestor, share the same pattern 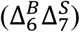, which is opposite to the one exhibited by *i*BSCc, *i.e. B. aphidicola* losses the capability to synthesize 8AONN and retains the capability to produce biotin from this precursor, whereas *S. symbiotica* exhibit the complementary pattern. Similar results are found when considering the reduction of each pathway individually, *i.e.* when only single pathway is considered redundant the GLRP include only the genes involved in the pathway (See Supplementary Figures S2-4 and Supplementary Table S6).

After the analysis of how the different GLRPs perform over the production rates of individual biomass components, the same study was conducted but using the meta-biomass production rate as a proxy to study the fitness of the alternative evolutionary scenarios. The simulation results, summarized in Fig 4, show that the value of the normalized meta-biomass production 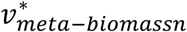 varies between 95 and 100%, compared to the putative ancestor (Fig 4a). Although this range is quite narrow (Fig 4b), it may still play a role in a selective process, since herein the solely stoichiometric rate is considered, and none other factor, such as the cost of protein synthesis, is considered. Moreover, in a first look the results indicate that the *i*BSCc GLRP 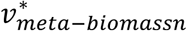is quite close to the one of the putative ancestor.

**Figure 4.**
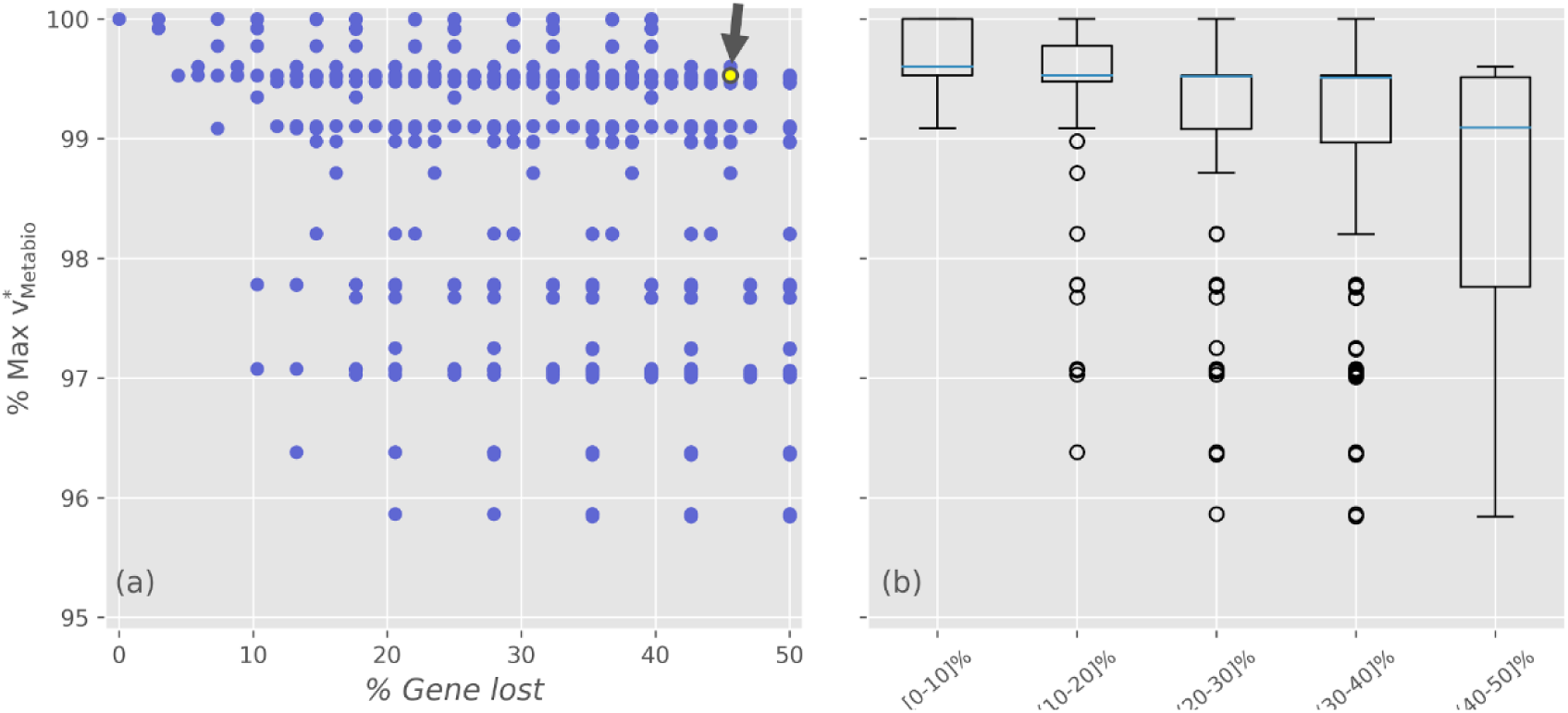
Optimal meta-biomass production rates for the different gene losses and retention scenarios. (a) Normalized optimal meta-biomass production rate plotted against the percentage of gene loss for each GLRP. (b) Different GLRPs were grouped by intervals of gene losses. The small arrow denotes the case of iBSCc, i.e. the cedar aphid consortium.

A deeper insight into the results shows among the minimal GLRPs, 48 from 128 of them (∼37%) also show a 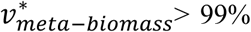. Furthermore, within this set it is possible to find opposite patterns, for example cases such as 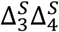 and 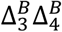 implies that it would be barely the same. Nevertheless, there are some particular patterns which are consistent, for example, all the minimal GLRP which include the loss 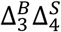 (*i.e.* the opposite pattern than the exhibited by *i*BSCc) imply 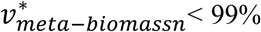. On the other hand, there are many GLRP which could represent disadvantages in terms of the systems growth rate. Such would be outcompeted by other more efficient organization of the complementation. Thus, considering only the structure of the networks of the endosymbionts, the simulation indicates that although many GLRP are possible, the one that is observed in nature, at least for the case of the cedar aphid consortium, has both large degree of gene loss and large growth rate.

## Discussion

### Small metabolic networks from bacterial endosymbionts

The reconstruction and metabolic analysis of *B. aphidicola* BCc and *S. symbiotica* SCc, coprimary endosymbionts of the cedar aphid *C. cedri*, has allowed, in first place, the revision of the annotation of these organisms’ genomes. Indeed, through the manual curation of these networks using the UM approach, it was possible to correct annotation errors in different enzymatic activities, and to identify previously unannotated metabolic genes. Moreover, the simulations performed here allowed the refinement of growth conditions and metabolic capabilities of these endosymbionts as compared to previous inferences from genomic analyses. Our study further confirms the validity of GEMs analysis for the phenotypic characterization of unculturable endosymbionts. On the other hand, the analysis of highly reduced metabolic networks, such as the case of *i*BCc98, bring into consideration methodological issues such as the case of the proton balance. Previously, a sensitivity analysis by Reed et al. (2003) in the genome-scale metabolic model of *E. coli* K12 *i*JR904 revealed that the net proton balance could be positive or negative depending on the carbon source used in the growth simulations, which would acidify or basify the environment, respectively. Alas, these predictions remain empirically untested.

The effect caused by proton balance in GEMs is generally low due to the size of these networks. However, in endosymbionts and other small networks, it may generate notorious consequences. In *i*BCc98, for instance, it considerably reduces the versatility of the metabolism of *B. aphidicola* BCc by coupling presumably independent processes, such as ATP synthesis and the folate cycle, and over-producing amino acids. Although this metabolic organization would be clearly disadvantageous for a free-living organism, for an endosymbiont member of a nutritional symbiosis it may be selected for at the level of the host. Indeed, a similar behaviour has been described recently as applied to the whitefly endosymbiont *P. aleyrodidarum* (Calle-Espinosa et al., 2016), where the growth of the organism is coupled with the overproduction of amino acids and carotenoids as a consequence of its low energetic capabilities. This phenomenon might play a relevant role in the evolution of nutritional endosymbiosis but it may also represent a methodological artefact as a consequence of the lack of knowledge on how to formulate in such a model the transport of protons through the membrane. One possibility would be the use of protons for the transport of compounds against their gradients. Although the scarcity of annotated transporters in the *B. aphidicola* BCc genome (Charles et al., 2011) does not seem to support this scenario, this problem falls within a more general umbrella, whereby the nature of the cell envelope (including both the membrane composition and the transport systems) of endosymbionts is largely unknown and might rely on contributions from the host (McCutcheon, 2016).

### Metabolic consequences of genome reduction

Simulations with and *i*BCc98 and *i*SCc236 indicate resemblance with previous metabolic analyses from endosymbionts and other bacteria with reduced genomes. We found that these two networks contain very few dispensable genes, with essentiality estimates being around 88% of genes for *i*SCc236, and 73% for *i*BCc98 (91% and 78%, respectively, according to MOMA). Although, these results may contradict the idea that the smaller the network, the higher the essentiality of its components, the percentage of essential genes predicted by *i*SCc236 drastically drops to ∼70%, when the cell wall genes are considered dispensable, and that the nucleotides and deoxynucleotides are provided by the host. Moreover, a further 15% and 5% of the genes, respectively, are genes that become essential after the deletion of another non-essential gene, thus displaying only a shallow degree of redundancy. Altogether, the amount of non-essential genes seems to positively correlate with the size of the network (Gil and Peretó, 2015). Moreover, a previous metabolic analysis of *B. aphidicola* APS showed that the network of this endosymbiont is also highly non-redundant, with 84% (94% according to MOMA) of genes being essential (Thomas et al., 2009). Although differences in the estimation algorithm and the selected criteria make these numbers not directly comparable, it does come to show that the *Buchnera* lineage evolved nearly minimal networks before the divergence of the Aphidinae and Lachninae aphid subfamilies, about 90 Mya. Moreover, although there are no available metabolic reconstructions for others *S. symbiotica* strains, the essentiality in the metabolic network of *S. symbiotica* SCc probably represents the high degree of genome reduction occurring in its symbiotic lineage, at 1.76 Mb and only 672 (Lamelas et al., 2011b). This is likely the result of more recent evolutionary processes, since *S. symbiotica* from hosts within the Aphidinae subfamily display genomes larger than 2.5 Mb and contain over 2000 CDSs (Burke and Moran, 2011; Foray et al., 2014; Manzano-Marín and Latorre, 2016). This might indicate that the genome reduction process in *S. symbiotica* SCc occurred after the divergence of the Lachninae subfamily, ca. 55 Mya. Two other genomes from obligate *S. symbiotica* strains have been described within this subfamily, obtained from *Tuberolachnus salignus* (tribe Tuberolachnini) and *Cinara tujafilina* (Eulachnini). The former contains a genome of only 650 kb and 495 CDSs (Manzano-Marín et al., 2016), while the genome of the latter is in an early stage of reduction at 2.5 Mb and 1602 CDSs (Manzano-Marín and Latorre, 2014). If the transmission of *Serratia* was vertical and no replacement occurred in the *C. tujafilina* clade, as it has been suggested (Manzano-Marín and Latorre, 2016), extreme reductive processes observed in the *i*SCc236 network may have been even more recent, possibly occurring no longer than 40 Mya.

Altogether, the two metabolic networks involved in the *C. cedri* consortium are highly constrained and fragile. This is also shown from the list of metabolic requirements that these organisms exhibit, which is increased by the high number of full and partial pathways that have been lost in both members of the consortium. Moreover, they show a high degree of integration, where both members have suffered massive losses, presumably due to division of labour with the bacterial partner. Cases of such losses are, for instance, the *B. aphidicola* BCc loss of the ability to produce cofactors like siroheme, biotin or THF, and, in *S. symbiotica* SCc, of the ability to produce several amino acids such as phenylalanine, threonine and branched amino acids. The model *i*BSCc establishes three cases of pathway sharing between the two symbionts, namely the biosynthesis of tryptophan, biotin and THF. These three pathways are partitioned between the two bacteria at the level of specific metabolites: anthranilate, 8AONN and shikimate, respectively (see Fig. 2). Those three exchanged metabolites are among the most permeable ones of the participant intermediates, as it was predicted by our previous chemoinformatic analysis of metabolic complementation (Mori et al., 2016). Finally, the endosymbiont metabolic networks also show the need for metabolic complementation from compounds synthesised by the host. This is reflected by the requirement for the incorporation of metabolic intermediaries, not just end-products, from the external compartment. Simulations with the metabolic network of *S. symbiotica* SCc show that two biosynthetic pathways are completed by the host. The loss of the first activities in the biosynthesis of siroheme and coenzyme A generate the need for the import of metabolic intermediaries. For example, a similar case has been previously observed in *Blattabacterium*, symbiont of the cockroach *Blattella germanica*, where the initial steps in the biosynthesis of terpenes have been lost (Ponce-de-Leon et al., 2013). These events are likely to commonly evolve in organisms under genome reduction processes, enabled by key factors such as the redundancy of pathways and the feasibility for transport due to permeability of the compound or to the existence or exaptation of generalist transporters.

### Emergence of metabolic complementation in endosymbiotic consortia

The simulations performed in this study with the reconstructed model of the consortium, *i*BSCc, reproduced the cases of metabolic complementation between the two bacterial partners that have been described in the literature, and predicted an additional one, the one involving the synthesis of THF in *S. symbiotica* SCc from the shikimate provided by *B. aphidicola* BCc. The emergence of metabolic complementation is a complex phenomenon and is not well understood from a theoretical perspective. Tryptophan biosynthesis is highly regulated at the transcriptional, translational and posttranslational levels. Notably, this pathway is negatively regulated through attenuation of the transcription of anthranilate synthase by tryptophan (Crawford, 1989), a process that has been suggested to facilitate the emergence of metabolic complementation (Mori et al., 2016). Genes for tryptophan synthesis from chorismate are present in most of the sequenced *B. aphidicola* genomes. In the cedar aphid, the genes encoding this pathway are divided in a *B. aphidicola* BCc plasmid containing *trpEG*, and a *S. symbiotica* SCc operon containing *trpABCD*. A strikingly identical case has been found to have occurred convergently in separate lineages of *Buchnera* and *Serratia* in the aphid *T. salignus* (Manzano-Marín et al., 2016). Moreover, the same complementation has been described in a completely different system, the one formed by *Ca.* Carsonella ruddii and secondary symbionts related to *Sodalis* and *Moranella*, in psyllids (Sloan and Moran, 2012). More complex cases of complementation in tryptophan biosynthesis also exist, such as the case of the mealybug symbionts, where this pathway seems to require the transport of multiple intermediaries, possibly including anthranilate (McCutcheon and von Dohlen, 2011).

Under this perspective, our simulations of all alternative GLRPs of metabolic complementations applied to the three shared pathways yielded novel results. We constructed models representing corresponding to alternative GLRPs from a hypothetical ancestor containing intact pathways in both symbionts, and compared how well they performed in maximizing the production of tryptophan, THF, biotin and meta-biomass, assuming that BCc and SCc are in proportion 1:1 in all the explored scenarios. Surprisingly, we’ve found that for the case of the tryptophan, the GLRP exhibited in the cedar aphid consortium behaved nearly optimality, and represented a quasi-minimal design in doing so. These results show that, from a structural point of view the actual distribution or division of a metabolic pathway between two organisms can perform almost as well as their ancestor while using a smaller gene repertoire. Although, the cost of protein synthesis and genome replication cannot easily be integrated in this kind of analysis, it is expected that a reduction in such costs will improve the growth efficiency of an organism (Mori et. al 2016). On the other hand, the results indicate that the GLRP exhibited in the biotin and THF pathways is suboptimal, and there are other GLRPs which allow a greater flux using the same number of genes. However, is worth to note that the demand for these cofactors is probably much smaller than the demand for tryptophan, and thus the selective pressure for an efficient production of THF and biotin may be less stringent than the case of the amino acid. This situation may also reflect the limits of the simultaneous optimization of the diverse metabolic performances of a given network. In any case, the GLRP of the tryptophan biosynthesis exhibited by the consortium of the cedar aphid is convergent with the metabolic solution observed in the symbiotic consortium of the psyllid *Heteropsylla cubana* (Martínez-Cano et al., 2015).

On the other hand, the reduction experiment performed evaluating the meta-biomass equation indicate that the global GLRP exhibited by the cedar aphid consortium, is quasi-optimal in terms of yield, and nearly minimal in terms of gene number. Moreover, additional factors can influence the given structure, and confer more benefits. For instance, the fact that the first and the last steps of the pathway are performed in different compartments reduces the possibility that an accumulation of the end-product tryptophan would inhibit the first biosynthetic step by attenuation. This is despite the fact that the inhibition binding site for tryptophan is highly conserved (Mori et al., 2016), which might be due to constraints in the enzyme functional architecture. Besides the structure of the complementation, the pathway kinetics is also likely to be involved in the function and evolution of the metabolic complementation. However, the complexity associated to a kinetic model and the lack of experimentally-based parameters makes such a model implausible at a genome scale as of today. Future efforts to model complete metabolic systems at a genome scale beyond stoichiometric constraints, by adding reaction kinetics and higher-level processes, such as the cost of protein production and turnover, will shed more light into the structure, function and evolution of metabolism, and in the emergence of metabolic complementation.

### Conclusions

The results and predictions obtained from GEMs, besides their intrinsic values, are useful as a tool to refine genomic and metabolic annotations. They also establish a powerful framework to interpret complex patterns of co-evolution, such as metabolic complementation. Here, we have reconstructed two genome-scale models from highly genome-reduced bacterial endosymbionts and integrate these models into a consortium model to study: (1) the requirements and exports of the bacterial partners to the host and to each other; (2) the robustness associated to reduced metabolic networks individually and by co-integration; and (3) the evolutionary constraints in the emergence of metabolic complementation designs. We could corroborate previously suggested scenarios for metabolic capabilities based on comparative genomic analyses. We also established that the cedar aphid consortium is composed not only of individual highly-reduced symbionts, but also that it is not far from a complete loss of metabolic redundancy and flexibility, thus making it a highly fragile partnership. Finally, we also showed that the patterns of metabolic complementation in this consortium are nearly minimal, in terms of gene content, and exhibit an almost optimal growth rate, and tryptophan production, with respect to a putative ancestor where the complemented pathways are still completely codded by each symbiont. Therefore, our results suggest a higher role of adaptive evolution in the emergence of metabolic complementation than previously thought, and more studies in different consortia with both similar and different patterns of complementation designs will be invaluable to confirm the generality of these conclusions.

## Author contributions

MP, FM, and JP conceived the work. MP, DT, and JCE developed the models. MP performed the simulations and analysed the data. All the authors discussed the results and wrote the manuscript.

## Acknowledgements

We would like to thank the Obra Social Programme of La Caixa Savings Bank for the doctoral fellowship granted to JCE. Financial support from Spanish Government (grant reference: BFU2015-64322-C2-1-R co-financed by FEDER funds and Ministerio de Economía y Competitividad) and Generalitat Valenciana (grant reference: PROMETEOII/2014/065) is grateful acknowledged. MM acknowledges financial support from the Simons Foundation. DT acknowledges support by a European Union grant from the Marie Curie ITN SYMBIOMICS (264774) and a grant from the Knut and Alice Wallenberg Foundation (2012.0075), given to Björn Andersson (Karolinska Institute) and Siv Andersson (Uppsala University).

## Supplementary Material

**Supplementary Text S1. Extended results.** The text include four sections, two including details about the reconstruction and metabolic capabilities of the endosymbiont individual models. The other two sections, correspond to the estimations of the proteome composition of *C. cedri* and to the construction of the meta-biomass.

**Supplementary Text S2. Extended Materials and Methods.** The text include details on the annotated genomes, constraint-based method employed, as well as the reconstruction of the plausible ancestor model of *C. cedri* used as a references point in the reduction experiments.

**Supplementary Figure S1. Conversion map predicted by *i*BCc98 and *i*SCc236.** Rows represent imported compounds, and columns, biomass components synthesized by each organism. Coloured squares in the same columns indicate the set of compounds required for the biosynthesis of a component. Purple and blue squares correspond to capabilities predicted by *i*BCc98 and *i*SCc236, respectively. Marked columns (*) correspond to compounds produced by both bacteria.

**Supplementary Figure S2. Optimal production rates of tryptophan for the reduced genes loss and retention experiments.** Normalized optimal production rate of tryptophan normalized with respect the optimal value exhibited. The small arrow denote the case of *i*BSCc, *i.e.* the cedar aphid consortium.

**Supplementary Figure S3. Optimal production rates of tetrahydrofolate for the reduced genes loss and retention experiments.** Normalized optimal production rate of tetrahydrofolate normalized with respect the optimal value exhibited. The small arrow denote the case of *i*BSCc, *i.e.* the cedar aphid consortium.

**Supplementary Figure S4. Optimal production rates of biotin for the reduced genes loss and retention experiments.** Normalized optimal production rate of biotin normalized with respect the optimal value exhibited. The small arrow denote the case of *i*BSCc, *i.e.* the cedar aphid consortium.

**Supplementary Table S1. Genome-scale metabolic model of *Serratia symbiotica* SCc *iS*Cc236.** The spreadsheet in XLS format includes the iSCc236 model description. It contains three tables: one that corresponds to the reactions; the second includes exchange fluxes; and in the third is the information of the metabolites.

**Supplementary Table S2. Genome-scale metabolic model of *Buchnera aphidicola* BCc *i*BCc98.** The file corresponds to the representation of the model iBCc98 in spreadsheet format, and includes the following four sheets: i) the list of reactions; (ii) listing of exchange fluxes; (iii) listing of metabolites; iv) the list of reactions and metabolic genes that were not included in the model.

**Supplementary Table S3. Genome-scale metabolic model of endosymbiotic consortium of the cedar aphid *C. cedri i*BSCc.** The file corresponds to the representation of the model *i*BSCc in spreadsheet format, and includes the following fiver sheets: i) the list of reactions; (ii) listing of exchange fluxes; (iii) listing of metabolites; iv) the optimal flux distribution that maximizes the growth of the whole system; and v) the exchange patterns between the consortium members.

**Supplementary Table S4. Results from the robustness analysis**. The table include two sheets. The first sheet list the results from the single KO *in-silico* experiments, whereas the second sheet include the results of the double KO *in-silico* experiments.

**Supplementary Table S5. Results from *in-silico* reduction experiments.** The table includes two sheets. The first sheet summarize the results from the reduction experiments and the second sheet includes a table with a mapping between Enzyme Subsets and the reactions IDs.

**Supplementary Table S6. Results from the reduced *in-silico* reduction experiments.** The table includes three sheets. Each sheet include the results of the reduction experiment for the tryptophan, tetrahydrofolate and biotin, when only the genes involved in each single pathway are considered for the experiment.

**Supplementary File S1. SBML models: iBCc98, iSCc236 and iBSCc.**

